# Structural basis for loading of Transcription Repair-Coupling factor Mfd onto stalled elongation complexes

**DOI:** 10.1101/2025.09.05.674597

**Authors:** Joshua Brewer, Eliza Llewellyn, James Chen, Elizabeth A. Campbell, Seth A. Darst

**Affiliations:** Laboratory of Molecular Biophysics, The Rockefeller University, New York, NY 10065; Laboratory of Molecular Pathogenesis, The Rockefeller University, New York, NY 10065

## Abstract

Transcription-coupled repair (TCR) is a nucleotide excision repair sub-pathway that preferentially removes lesions from the DNA template-strand stalling RNA polymerase (RNAP) elongation complexes (ECs). In bacteria, the superfamily 2 Mfd translocase mediates TCR by displacing stalled ECs and recruiting Uvr(A)B. Using cryo-electron microscopy, we previously visualized seven Mfd-EC complexes spanning the ATP-dependent Mfd loading and EC displacement pathway [L1 -> L2(ADP) -> C1(ATP) -> C2(ATP) -> C3(ADP) -> C4(ADP) -> C5(ATP)]. The first intermediate (L1) was poorly resolved (4.1 Å nominal resolution) due to low particle occupancy. The pathway is characterized by very large Mfd structural transitions, notably the L1 -> L2 transition. Here, we pre-loaded Mfd with ATP in the presence of the *γ*-phosphate mimic, BeF_3_^−^, limiting rounds of ATP hydrolysis. The resulting accumulation of early intermediates allowed us to resolve the L1 intermediate to 3.5 Å nominal resolution, revealing bound ADP-BeF_3_^−^. We also identified a new intermediate between L1 and L2, L1.5, providing further insight into Mfd conformational changes during loading.

**Graphical Abstract:** 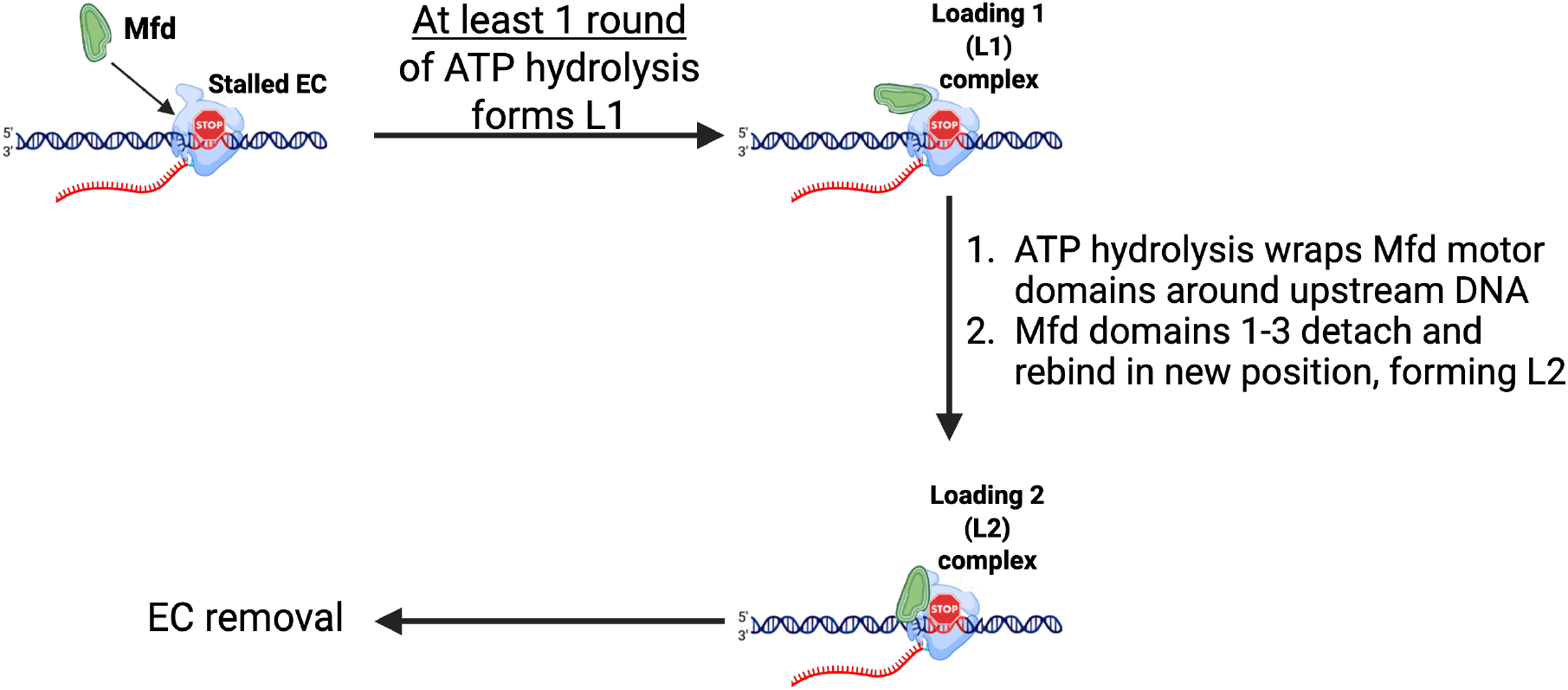

## Introduction

DNA template strand (t-strand) lesions that block elongation by RNA polymerase (RNAP) are targeted for preferential repair in a process called transcription-coupled repair (TCR) [1–3]. UV-induced cyclobutane dimers are the most common lesions that induce TCR [4,5]. In TCR, the stalled RNAP elongation complex (EC) serves as an efficient detector of t-strand lesions that then becomes a privileged entry point into the nucleotide excision repair (NER) pathway, mediated by a transcription repair coupling factor (TRCF). The superfamily 2 (SF2) Mfd translocase [6] has been shown *in vitro* and *in vivo* to be a bacterial TRCF [7–11].

In its apo form, Mfd is autoinhibited, inactive in binding to or translocating along DNA, binding to NER factor UvrA, or hydrolyzing ATP [6,12,13]. This is attributed to the compact inactive conformation of apo-Mfd, where the UvrA-interacting module is occluded by Mfd domain 7 [D7; [14]]. Stable association of Mfd with the EC requires many rounds of ATP hydrolysis [15].

After engaging with the EC, Mfd translocase activity can overwind the upstream region of the EC transcription bubble, facilitating displacement of the RNA transcript and transcription bubble reannealing [6,12,15–21]. After disruption of the EC, the Mfd-RNAP complex remains on the DNA and continues to slowly translocate in the downstream direction processively over thousands of base pairs [18,22,23]. The Mfd-RNAP complex rapidly dissociates from the DNA once it is intercepted by UvrA_2_B [24,25].

Previously, we used cryo-electron microscopy (cryo-EM) to visualize the ATP-dependent pathway for Mfd engaging with and attempting to displace the EC [15]. We visualized seven distinct Mfd-EC complexes in both ATP and ADP-bound states spanning the Mfd loading and EC displacement pathway:

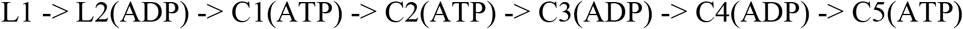

In this earlier study, the first Mfd-EC loading intermediate (L1) was poorly populated, yielding a relatively low-resolution structure (4.1 Å nominal resolution) in which the nucleotide occupancy of the Mfd active site was unclear. The pathway is characterized by very large Mfd structural transitions, most notably the L1 -> L2 transition, where the RecG-like SF2 translocase domains (TDs) translocate at least 7 base pairs along the upstream duplex DNA of the EC, rotating ∼258° about the DNA and undergoing an overall translation of ∼63 Å while hydrolyzing at least 7 molecules of ATP. Other domains of Mfd undergo similarly large motions. The rotation of the Mfd TDs around the DNA, combined with rearrangements of other Mfd domains, result in a conformation in which Mfd is topologically wrapped around the upstream duplex DNA. This persistently ‘wrapped’ arrangement of the Mfd-RNAP complex was proposed to be the source of its high processivity on DNA [15,18,22,23]. While the endpoints of the initial structural transition (L1 and L2) were resolved, the detailed pathway of Mfd domain motions during the transition was unclear.

Here, we pre-loaded Mfd with ATP in the presence of BeF_3_^−^, allowing Mfd to undergo a limited number of ATP hydrolysis cycles before being trapped by BeF_3_^−^ binding prior to ADP release. The resulting accumulation of particles in the early steps of the pathway allowed us to resolve the L1 intermediate to 3.5 Å nominal resolution, revealing nucleotide occupancy. We also identified an additional loading intermediate between L1 and L2 (L1.5) that clarifies the rearrangements undertaken by Mfd during loading.

## Materials and Methods

### Key Resource Table

[].

### Lead Contact and Materials Availability

All unique/stable reagents generated in this study are available without restriction from the Lead Contact, Seth A. Darst.

### Experimental Model and Subject Details

RNAP core (α_2_ββ’ω) and Mfd are proteins found in *Eco*. For protein expression, *Eco* BL21(DE3) [*Eco* str. B F^−^ *ompT gal dcm lon hsdS*_*B*_(*r*_*B*_^*–*^*m*_*B*_^*–*^) λ(DE3 [*lacI lacUV5*-*T7p07 ind1 sam7 nin5*]) [*malB*^+^]_K-12_(λ^S^)] was used.

### Method Details

Structural biology software was accessed through the SBGrid consortium [26]. No statistical methods were used to predetermine sample size. The experiments were not randomized. The investigators were not blinded to allocation during experiments and outcome assessment.

### Protein expression and purification

*Eco* core RNAP and *σ*^70^ were separately overexpressed and purified as previously described [15,27]. *Eco* Mfd was overexpressed and purified as previously described [15].

### Mfd-EC Displacement Assay

50 picomoles of forward primer (5’ cggaacattacgaacgatgg 3’ IDT), used later for PCR amplification of pAR1707 [28], was combined with 50 picomoles of *γ*-^32^P-ATP, 2 µL of T4 polynucleotide kinase (10,000 U/mL), and T4 buffer (New England Biolabs). Radiolabeling reactions ran for 45 min at 37°C. After a 20-minute heat treatment at 65°C, radiolabeled primers were purified using a NucAway spin column (Invitrogen AM10070). PCR reactions were performed with radiolabeled forward primers and unlabeled reverse primers from pAR1707 template (Thermo Scientific Phusion High-Fidelity DNA polymerase F-530XL). Amplicons were run on an SDS polyacrylamide gel and the purified band was cut out and dissolved in Crush/Soak elution buffer (1 M Na-acetate, pH 5.5, 1 mM EDTA) before being spin filtered in an Ultra-free MC tube (Millipore UFC30VV00) and precipitated in 100% ethanol, washed with 70% ethanol, and finally resuspended in RNase free 10 mM Tris-HCl, pH 8.0 (Invitrogen AM9855G).

*Eco* RNAP holoenzyme (E*σ*^70^) was reconstituted by incubating core RNAP (E) (10 nM final) with σ^70^ (50 nM final) at 37°C for 15 min. E*σ*^70^ was incubated with radiolabeled DNA fragments (0.4 nM) for 10 minutes at 37°C to form transcription initiation complexes. Transcription was then initiated by adding ApU (200 uM), ATP (2 mM), CTP (50 uM), GTP (50 uM), 3’-deoxy-UTP chain terminator (100 *μ*M) (all NTPs: TriLink Biotechnologies) and heparin (10 *μ*g/ml; Sigma-Aldrich H3149) at 37°C for 10 min in transcription buffer (100 mM Tris-HCl, pH 8.0, 500 mM KCl, 100 mM MgCl_2_, 1 mM EDTA, 10 mM dithiothreitol, 50 μg/mL BSA). In conditions where a non-hydrolyzable NTP analog was used to halt Mfd activity, either 20 mM Na-orthovanadate (VO_4_^3−^; Sigma-Aldrich 450243), 13.95 mM Aluminum trifluoride (AlF_3_; Sigma-Aldrich 449268), or 25 mM Beryllium trifluoride (BeF_3_^1−^; Sigma-Aldrich AA1610414) were added to the transcription reaction simultaneously with other NTPs and heparin. Finally, Mfd (250 nM) was added to the reaction and time points were taken by adding samples directly to the running gel. Native gels were cast using 4.5% polyacrylamide [(37.5:1) acrylamide/bis-acrylamide (2.7% crosslinker), 0.15% TEMED (Thermo Scientific 17919), and 1% APS (Sigma-Aldrich A3678). Gels were run at 4°C at 24 mA for approximately 19 hours. Gels were mounted onto Whatman filter paper and placed into a gel dryer for 30 min at 40°C. Gels were exposed to a phosphor screen for 6 hrs at 4°C and imaged using a Typhoon (Cytiva). Band intensities were quantified using ImageJ. Stall intensity % was calculated as follows: (Stall band intensity)/(Stall band intensity + DNA band intensity) using background-subtracted values.

### Preparation of Mfd-EC complexes for cryo-EM

*Eco* core RNAP (0.5 mL of 5 mg/mL protein) was injected into a 10/300 Superose 6 Increase column (Cytiva) equilibrated with 10 mM Tris-HCl, pH 8.0, 100 mM KCl, 5 mM MgCl_2_, and 2.5 mM dithiothreitol (DTT). The peak fractions of the eluted protein were concentrated by centrifugal filtration (EMD-Millipore-30K MWCO) to 25 µM protein concentration. The above steps were also performed for purified Mfd. ECs were reconstituted by hybridizing RNA with t-strand DNA (stepwise decreasing temperature from 95°C) before adding *E. coli* RNAP (RNAP:DNA ratio of 1:1.1) at room temperature for 15 minutes. Nt-strand DNA was finally added for 10 minutes at room temperature. ATP (2 mM), BeF_3_^−^ (25 mM), 24.872 uM Mfd, and 12.422 uM EC (final concentrations) were all combined in Cryo-EM buffer (150 mM KCl, 20 mM Tris pH 8.0, 10 mM MgCl_2_, 10 mM DTT). The sample was incubated for 15 min at 37°C, then 3-([3-cholamidopropyl]dimethylammonio)-2-hydroxy-1-propanesulfonate (CHAPSO) was added to a final concentration of 8 mM [29], and the sample was kept at room temperature prior to grid preparation.

### Cryo-EM grid preparation

C-flat holey carbon grids (CF-1.2/1.3-4Au; Protochips) were glow-discharged for 20 s before the application of 3.5 μL of the sample described above. After blotting for 3–4.5 s, the grids were plunge-frozen in liquid ethane using an FEI Vitrobot Mark IV (FEI) with 100% chamber humidity at 37°C.

### Cryo-EM data acquisition and processing

Grids were imaged using a 300-keV Titan Krios (FEI) equipped with a K3 Summit direct electron detector (Gatan). Images were recorded with Leginon [30] in counting mode with a pixel size of 1.0825 Å (our 2^nd^ dataset was collected with a pixel size of 1.03Å and subsequently binned to 1.0825Å) and a defocus range of -1 to -2.5 μm. Data were collected with a dose rate of 28 *e*^-^ per Å^2^ per s. Images were recorded over a 2 s exposure with 0.05 s frames (40 total frames) to give a total dose of 51.2 *e*^-^/Å^2^ (2^nd^ dataset had a total dose of 68.2 *e*^-^/Å^2^). Dose-fractionated videos were gain-normalized, drift-corrected, summed, and dose-weighted using MotionCor2 [31]. The contrast transfer function (CTF) was estimated for each summed image using the Patch CTF module in cryoSPARC2 (CS2) [32]. Particles were picked and extracted from the dose-weighted images with a box size of 256 px using CS2 Blob Picker and Particle Extraction.

Coordinates pointing to particles of ice were extracted as faux particles and used to generate an initial decoy 3D model in CS2 (ab initio reconstruction) in order to remove junk particles from initial particle stacks during the ‘decoy’ procedure. Multiple rounds of CS2 Hetero Refinement of all blob-picked particles used multiple 3D decoys along with an *Eco* core RNAP 3D template (PDB 6ALH) with all nucleic acids removed (low pass filtered to 60 Å resolution), to perform initial de-junking. The original purpose for the 2^nd^ data collection was to identify more particles of L1.5 to improve map resolution; however, no additional L1.5 particles could be successfully identified, whereas additional L1 particles were identified. Both L1 and L1.5 were classified from their respective parent particle stacks via focused 3D classification in Relion3 [33]. Both classes were subjected to two rounds of successive Bayesian Polishing in Relion3. Finally, CS2 non-uniform (NU) refinements were performed for each resulting class, yielding two different structures: L1 (96,588 p, 3.5 Å nominal resolution) and L1.5 (13,101 p, 4.3 Å nominal resolution).

The heatmap distribution of particle orientations and half-map FSCs were calculated using CS2. 3D Fourier shell correlation (3dFSC) calculations were performed using 3DFSC [34]. Local-resolution calculations were performed using blocres and maps were locally filtered using blocfilt (Bsoft package) [35].

### Model building and refinement

*L1*. The initial model for L1 was derived from PDB 6X26 [15]. The model was manually fit into the cryo-EM density maps using ChimeraX [36] and rigid-body and real-space refined using PHENIX real-space-refine [37,38]. For real-space refinement, rigid-body refinement was followed by all-atom and *B* factor refinement with Ramachandran and secondary structure restraints. Models were inspected and modified using COOT [39].

*L1*.*5*. The initial model for L1.5 was derived from PDB 6X50 [40].

### Data and code availability

The cryo-EM density maps have been deposited in the EMDataBank under accession codes EMD-48776 [L1(ADP-BeF_3_)] and EMD-48802 [L1.5]. The atomic coordinates have been deposited in the Protein Data Bank under accession codes 9N07 [L1(ADP-BeF_3_)] and 9N11 (L1.5).

## Results

### Mfd activity is sensitive to *γ*-phosphate mimics

ATP hydrolysis is required for Mfd loading to achieve a stable Mfd-EC complex (Kang et al. 2021). For example, incubating a stalled EC with transition state analog ADP + vanadate (VO_4_^3−^) [41] does not support Mfd-EC complex formation, but ATP + VO_4_^3−^ does [15]. Although ATP + VO_4_^3−^ supports Mfd-EC complex formation, it does not support EC displacement, indicating that multiple rounds of ATP hydrolysis are required for EC displacement [15]. Similarly, ATP with ground state analogs AlF_3_^−^ or BeF_3_^−^ also do not support EC displacement (Figures 1b and 1c, Figure 1-figure supplement 1). Thus, pre-loading Mfd with ATP in the presence of *γ*-phosphate mimics VO_4_^3−^, AlF_3_^−^, or BeF_3_^−^ allows Mfd to undergo a limited number of ATP hydrolysis cycles, which are obligate for Mfd-EC complex formation, but not enough cycles to complete EC displacement.

**Figure 1.**
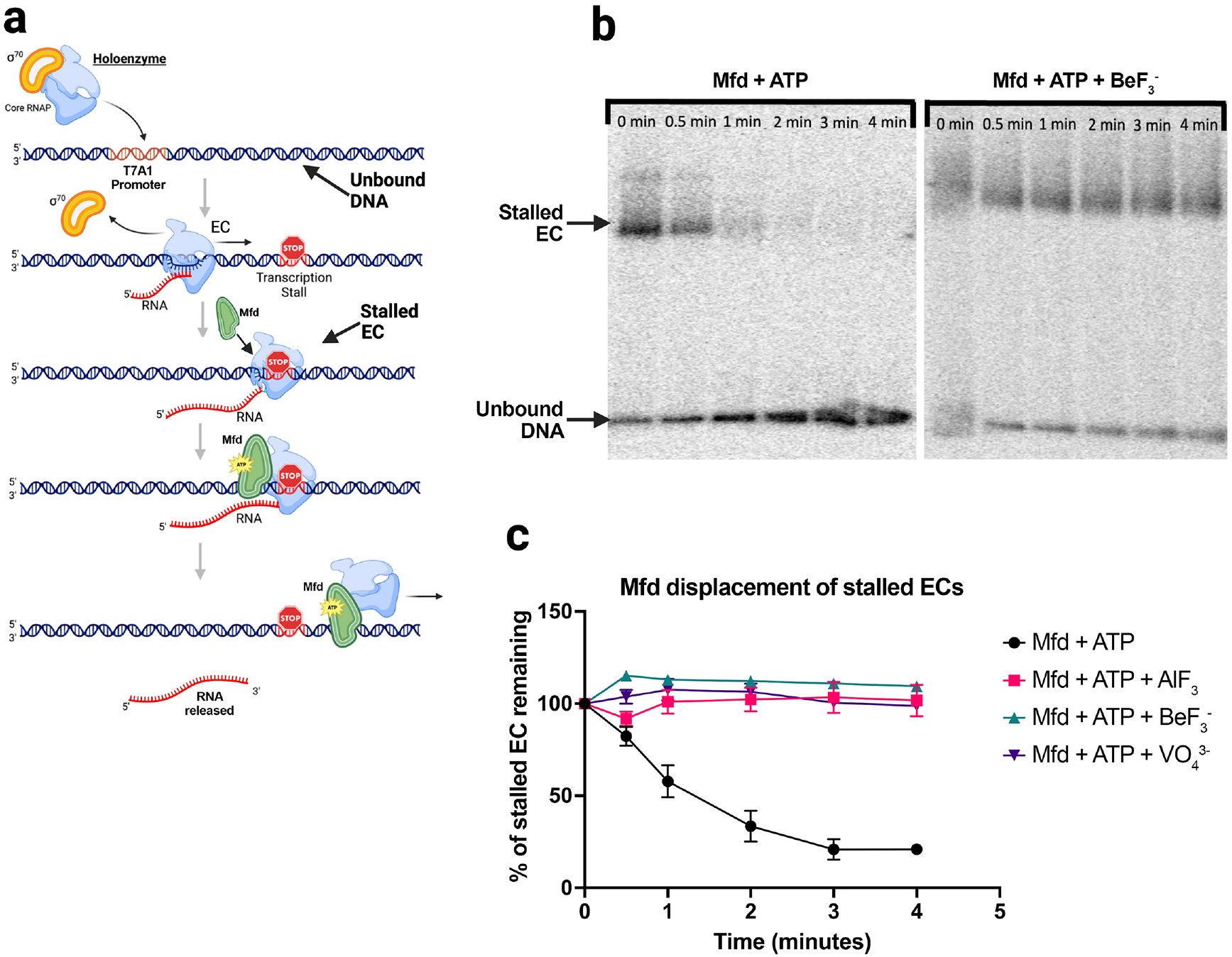
Non-hydrolyzable ATP analogs inhibit Mfd-mediated displacement of stalled ECs. (**a**) Schematic depicting native gel-shift experiments used to monitor Mfd-mediated displacement of stalled ECs. *Eco* holoenzyme was added to a radiolabeled linear dsDNA fragment containing the strong T7A1 promoter (Figure 1-figure supplement 1a). Stalled ECs were formed via the introduction of dinucleotide primter (ApU, 200 uM final concentration), a subset of NTP substrates (ATP, 2 mM: CTP, 50 uM; GTP, 50 uM) and a chain terminator (3’-deoxy-UTP, 100 *μ*M). The *γ*-phosphate mimics Sample-dependent non-hydrolyzable ATP-analog was introduced prior to Mfd [NaVO_4_^3−^ (20 mM final concentration), or AlF_3_ (14 mM), or BeF_3_^−^ (25 mM)]. Mfd hydrolyzes ATP to displace, translocate, and ultimately dissociate the stalled EC from the DNA fragment. (**b**) The left panel shows the effect of Mfd-mediated removal of the stalled EC via native electrophoretic mobility shift analysis; after 1 to 2 minutes of incubation with Mfd and ATP, the low mobility EC (‘stalled complex’) is disrupted. The right panel shows the inhibitory effect of BeF_3_^−^. (**c**) Quantification of Mfd-mediated removal of stalled ECs from the linear DNA fragment and the potent inhibitory effects of all three tested *γ*-phosphate mimics. Error bars denote standard deviation of n=3 measurements. Representative gels for AlF_3_ and VO_4_^3−^ analyses are shown in Figure 1-figure supplement 1b and 1c).

### Two structures early in the Mfd loading pathway

We reasoned that limiting the number of ATP hydrolysis cycles that Mfd could complete by using ATP and a *γ*-phosphate mimic would enrich Mfd-EC complexes early in the loading pathway. We incubated stalled ECs with Mfd pre-loaded with ATP [Mfd(ATP)] in the presence of 25 mM BeF_3_^−^ (Figures 1b,1c, and 2a), then analyzed the resulting complexes by single particle cryo-EM (Figure 2, Figure 2-figure supplement 1). Steps of maximum-likelihood classification [42] revealed two predominant Mfd-EC structures. As expected, the sample was highly enriched in L1 (88% of the particles; Figure 2b, Figure 2-figure supplement 1). Unexpectedly, the remainder of the particles (12%) defined a new structural class between L1 and L2 that we term L1.5 (Figure 2c, Figure 2-figure supplement 1).

**Figure 2.**
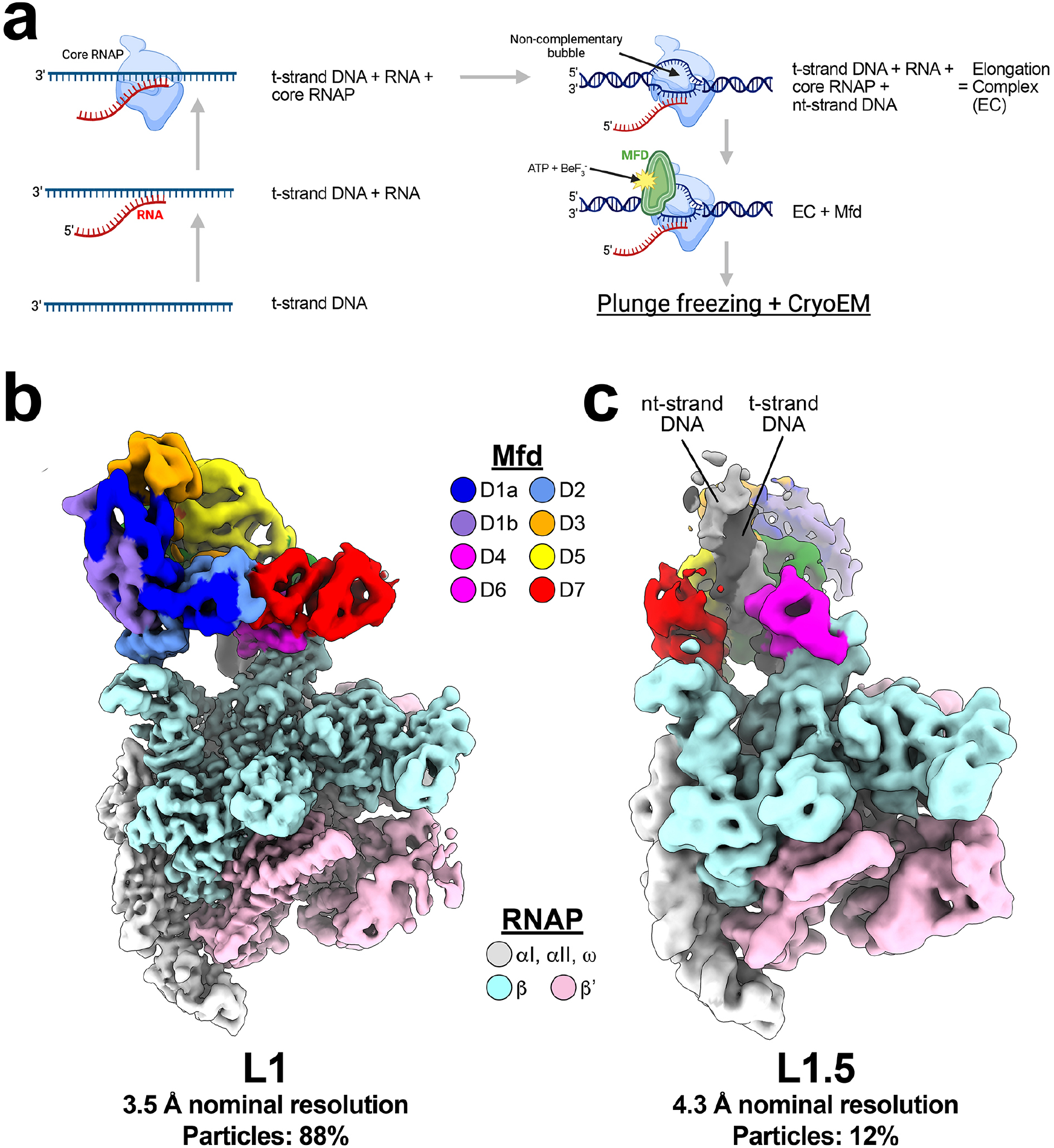
Two structure early in the Mfd loading pathway. (**a**) Schematic for reconstituting the stalled EC employed in cryo-EM experiments. To assemble complexes enriched for early intermediates in the Mfd loading pathway, we annealed partially complementary RNA to t-strand DNA, added core RNAP, then finally nt-strand DNA [with partial complementarity to the t-strand, resulting in a bubble resistant to annealing [15]]. The subsequent addition of Mfd, ATP (2 mM final concentration), and BeF_3_^−^ (25 mM) allowed formation of early intermediates via limited rounds of ATP hydrolysis before BeF_3_^−^ can associate with Mfd to inhibit further activity. (**b**) Local-resolution filtered [35] cryo-EM density for L1 intermediate with associated resolution and % particle contribution to overall population observed. Color legend for domains of RNAP and Mfd shown. (**c**) Local-resolution filtered [35] cryo-EM density for L1.5 intermediate with associated resolution and % particle contribution to overall population observed. Color legend for domains of RNAP and Mfd shown. The upstream duplex DNA (t-strand and nt-strand) is labeled. The RNA transcript present in the compexes is not visible in these views.

### High-resolution L1 structure confirms ATP in the Mfd active site

L1 was proposed to be the first stable intermediate on the Mfd-EC loading pathway [15]. The nucleotide state of the Mfd active site was previously unresolved due to the poor resolution of the structure [4.1 Å nominal resolution; [15]]. By limiting the number of Mfd ATP hydrolysis cycles in our current study, the population of particles in the L1 state was enriched, giving rise to a much better resolved structure (3.5 Å nominal resolution; Figures 2b and 3, Figure 2-figure supplement 1). The previous L1 model (4.1 Å nominal resolution) and the current L1 model (3.5 Å resolution) are very similar in overall architecture (1.18 Å root-mean-square deviation over 3,731 *α*-carbons), but the new model clearly resolves ADP•BeF_3_^−^ in the Mfd active site (Figure 3). BeF_3_^−^ is a ground-state analog, suggesting that the L1 state contains Mfd(ATP) during the normal ATP hydrolysis condition.

**Figure 3.**
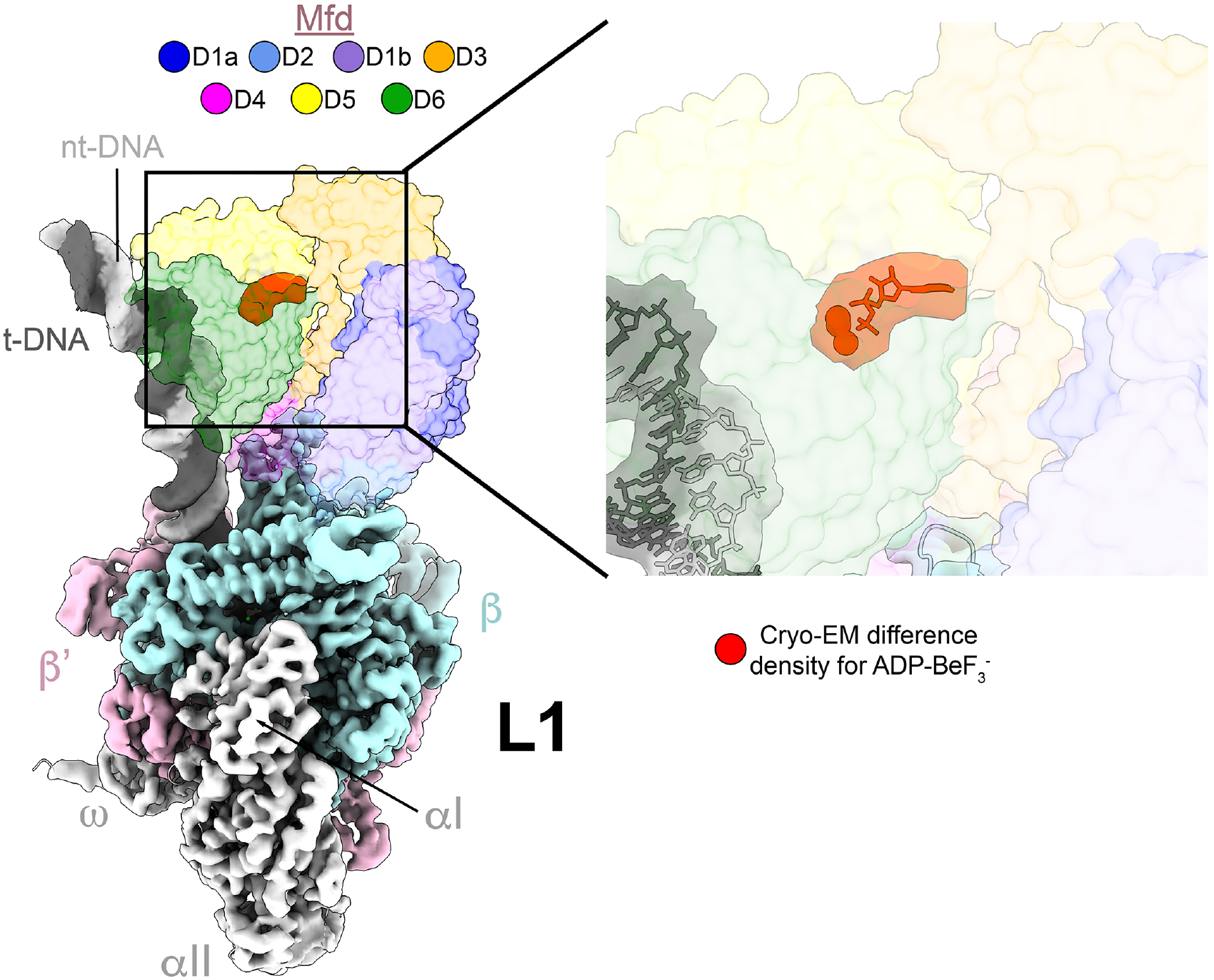
High resolution structure of L1 confirms presence of bound ADP-BeF_3_^−^ in Mfd active site. The L1 intermediate is shown on the left [local-resolution filtered cryo-EM map [35] for RNAP and nucleic acids, with transparent molecular surface of Mfd colored according to key (top). DNA stands and RNAP subunits are individually labelled and distinguished by color. Cryo-EM difference density (zoomed in rightmost panel) shows clear density corresponding to the presence of ADP-BeF_3_^−^ (shown in red with model fit into difference density).

### ATP loading of Mfd in conjunction with BeF_3_^−^ traps the Mfd-EC complex in a new intermediate

The ATP-dependent transition from L1 to L2 involves dramatic conformational changes in the positioning of Mfd domains. In L1, the Mfd TDs engage with the upstream duplex DNA of the EC far upstream, centered at about -30 with respect to the 3’-end of the RNA in the EC. In L2, the Mfd TDs are centered at about -21 on the upstream duplex DNA and rotated around the DNA about 258° [15]. Thus, in the L1->L2 transition, the Mfd TDs translocate on the upstream duplex DNA in the downstream direction (i.e. towards the RNAP) while the RNAP and the RNAP-bound Mfd-D4(RID) remain stationary. During this process, a long helix (the Relay Helix) connecting Mfd D4(RID) to D5(TD1) unfolds and wraps around the DNA, imparting the translocating Mfd-EC complex its striking processivity [15,24]. It was proposed in Kang et al. (2021) that the Mfd D1-D3 structural module disengages from the Mfd-EC complex, opening a path for the TDs to corkscrew around the DNA to reach their position in L2. D1-D3 was proposed to subsequently rebind in a different position, forming L2 [15,24].

In our new cryo-EM dataset, most particles (88%) have undergone one (or at most, a few) rounds of ATP hydrolysis, generating L1 (Figures 2b and 3). However, a smaller population of particles (12%) has successively hydrolyzed a limited number of ATP molecules prior to associating with BeF_3_^−^, moving beyond L1 but not yet achieving L2, revealing a new intermediate, L1.5 (Figures 2c and 4, Figure 2-figure supplement 1). In L1.5, the Mfd TDs are centered at about -25 and rotated around the DNA about 181° (with respect to L1), approximately half-way between L1 and L2 (Figure 4).

**Figure 4.**
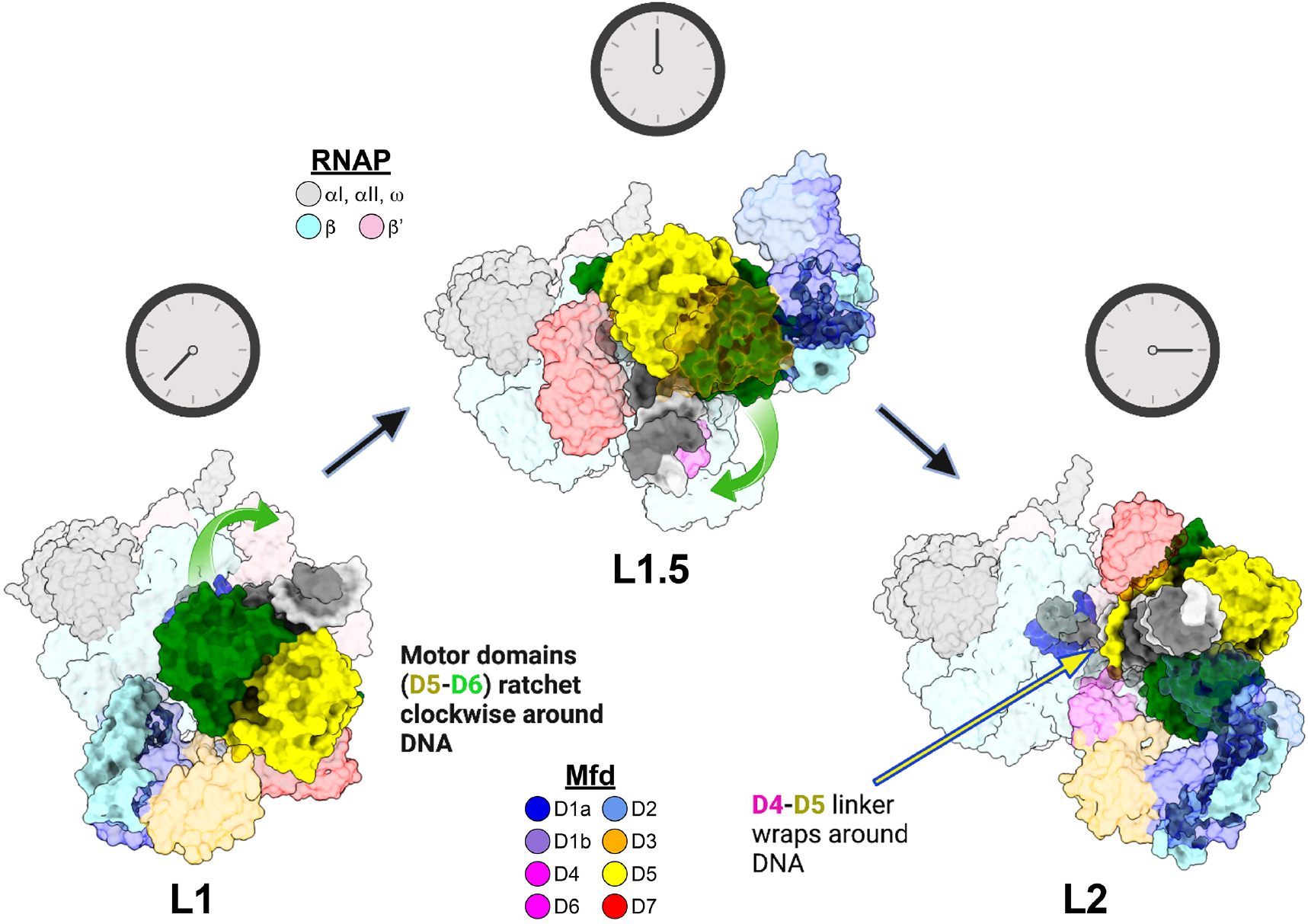
L1.5 structure illustrates pathway for transition from L1 to L2. Mfd-EC complexes L1 (left, this work), L1.5 (middle, this work), and L2 [right, 6X2F; [15]] illustrate rearrangements of Mfd structural elements to accomplish loading. The complexes are shown as molecular surfaces (RNAP subunits and Mfd domains color-coded as shown), aligned via the RNAP, and viewed from the upstream duplex DNA direction. The Mfd RecG-like motor domains (D5, yellow; D6, green] corkscrew around the upstream dsDNA in the clockwise direction while hydrolyzing ATP. The relay helix (D4-D5 Mfd linker, labeled in L2) gradually wraps around the upstream duplex DNA as motor domains circle around the DNA and the RNAP-Interacting-Domain [D4(RID)] remains attached to the RNAP (highlighted in rightmost complex shown). D1-D3 appear to temporarily dissociate from the other Mfd domains as motor domains circle the upstream duplex DNA and rebind in a different overall conformation (L1.5) that persists (with minor repositioning relative to the motor domains) until the final formation of L2 where Mfd has achieved loading.

## Discussion

In Kang et al. (2021), Mfd was visualized undergoing its complete ATP hydrolysis cycle, engaging with and attempting to disrupt an EC. The ECs were stalled by nucleotide deprivation on a nucleic acid scaffold containing a non-complementary transcription bubble that could not rewind (Figure 2 - figure supplement 1a). Thus, despite undergoing cycles of ATP hydrolysis, Mfd was unable to efficiently disrupt the ECs, facilitating the visualization of intermediates. Seven distinct Mfd-EC complexes were visualized and placed in a pathway, providing a structural basis for understanding the extensive remodeling of Mfd upon its engagement and disruption of the EC. The presence of ATP throughout the sample preparation allowed multiple rounds of ATP hydrolysis, driving the complexes through the pathway, explaining why the early intermediates (L1 and L2) were poorly populated; L1 and L2 contained 4% and 5% of the particles, respectively. The small number of particles gave rise to relatively low-resolution cryo-EM maps (L1, 4.1 Å nominal resolution; L2, 4.0 Å nominal resolution). As a result, the nucleotide occupancy of the Mfd active site in L1 was not able to be determined.

In this current study we populated complexes early in the pathway by incubating the stalled EC with Mfd, ATP, plus a *γ*-phosphate mimic, BeF_3_^−^. Following ATP hydrolysis by Mfd, BeF_3_^−^ can substitute for the leaving inorganic phosphate before ADP has been released [43,44], trapping the complex after one or only a few rounds of ATP hydrolysis. In this way, the sample was heavily shifted towards populating early complexes. L1 contained 88% of the particles, giving rise to a 3.5 Å reconstruction (Figures 2b and 3, Figure 2 - figure supplement 1). The better resolution of the L1 cryo-EM map revealed that ADP-BeF_3_^−^ occupied the Mfd active site (Figure 3), consistent with biochemical findings that ATP hydrolysis was required to reach L1 [15].

In addition to increasing the population of L1, a new complex between L1 and L2, L1.5, was populated (12% of the particles). In the presence of BeF_3_^−^, L2 and subsequent states were not populated at all (Figure 2 - figure supplement 1). In L1.5, the Mfd-TDs translocating (corkscrewing) on the upstream duplex DNA towards the RNAP were trapped approximately half-way between L1 and L2 (Figure 4), confirming and elaborating the proposed L1 -> L2 transition pathway [15].

The ability of cryo-EM to visualize mixtures of biological macromolecules in relatively native environments allows the analysis of complex assembly or activity pathways *in vitro*, revealing unprecedented insights [15,45–47]. By their nature, active biological pathways lead to extreme heterogeneity. In the case of NTPase-driven processes, the use of *γ*-phosphate mimics to allow limited rounds of ATP hydrolysis can reduce the heterogeneity and stabilize transient states that otherwise would not be populated, providing insight into NTP-hydrolysis driven pathways difficult to obtain in other ways.

## Supplementary data

Supplementary Data are available online.

## Acknowledgments

We thank members of the Darst and Campbell Laboratories for helpful discussions, and M. Ebrahim, J. Sotiris, and H. Ng at The Rockefeller University Evelyn Gruss Lipper Cryo-Electron Microscopy Resource Center for help with cryo-EM data collection and analysis. Some of the work reported here was conducted at the Simons Electron Microscopy Center (SEMC) and the National Resource for Automated Molecular Microscopy (NRAMM) located at the New York Structural Biology Center, supported by grants from the NIH National Institute of General Medical Sciences (P41 GM103310), NYSTAR, the Simons Foundation (SF349247), the NIH Common Fund Transformative High Resolution Cryo-Electron Microscopy program (U24 GM129539) and NY State Assembly Majority. This work was supported by NIH R35 GM118130 to S.A.D.

## Author contributions

Conceptualization; J.B., E.A.C., S.A.D. Cloning, protein purification, biochemistry: J.B., E.L. Cryo-EM specimen preparation, data collection, and processing: J.B., E.L., J.C. Model building and structural analysis: J.B., E.A.C., S.A.D. Funding acquisition and supervision: E.A.C., S.A.D. Manuscript first draft: J.B., S.A.D. All authors contributed to finalizing the written manuscript.

## Data Availability

The data underlying this article are available in the article and in its online supplementary material.

