## Supplemental Information for "Structural basis for loading of Transcription Repair-Coupling factor Mfd onto stalled elongation complexes"

### Supplemental Data

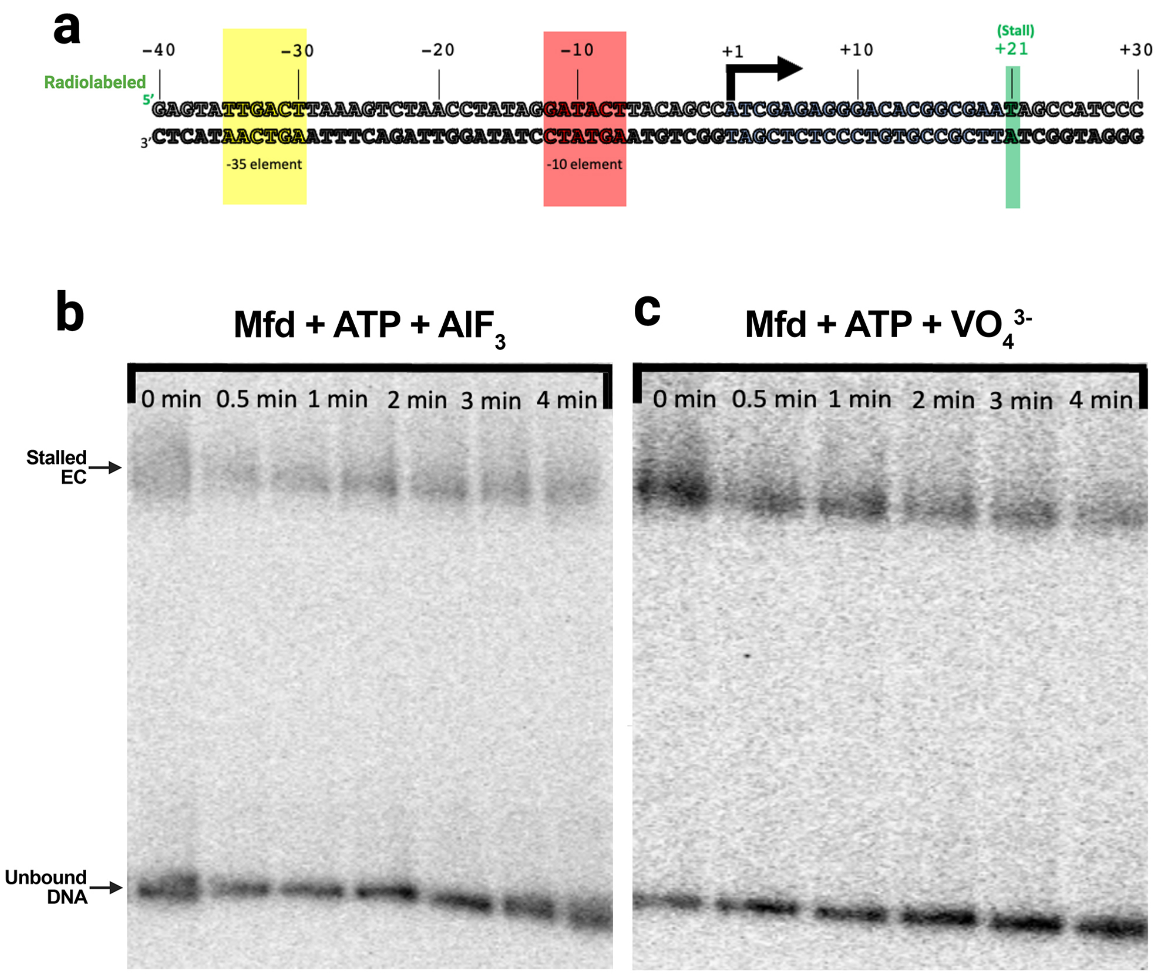

**Supplementary Figure 1.**

**(a)** Linear dsDNA T7A1 promoter fragment sequence used in native gel-shift experiments. Annotations include the core promoter elements (-35 element, yellow; -10 element, red), the transcription start site (+1, denoted with the bent arrow pointing in the direction of transcription), and the site of the EC stall (+21 shaded green, where RNAP is expected to incorporate chain-terminating 3'-deoxy-UTP). Position of g-$^{32}$P-ATP-labelled 5' end of the nt-strand is also shown in green.

**(b, c)** Representative native gel-shift results monitoring Mfd-mediated removal of stalled ECs in the presence of **(b)** $AlF_3$ (14 mM; left panel) or **(c)** $VO_4^{3-}$ (20 mM; right panel).

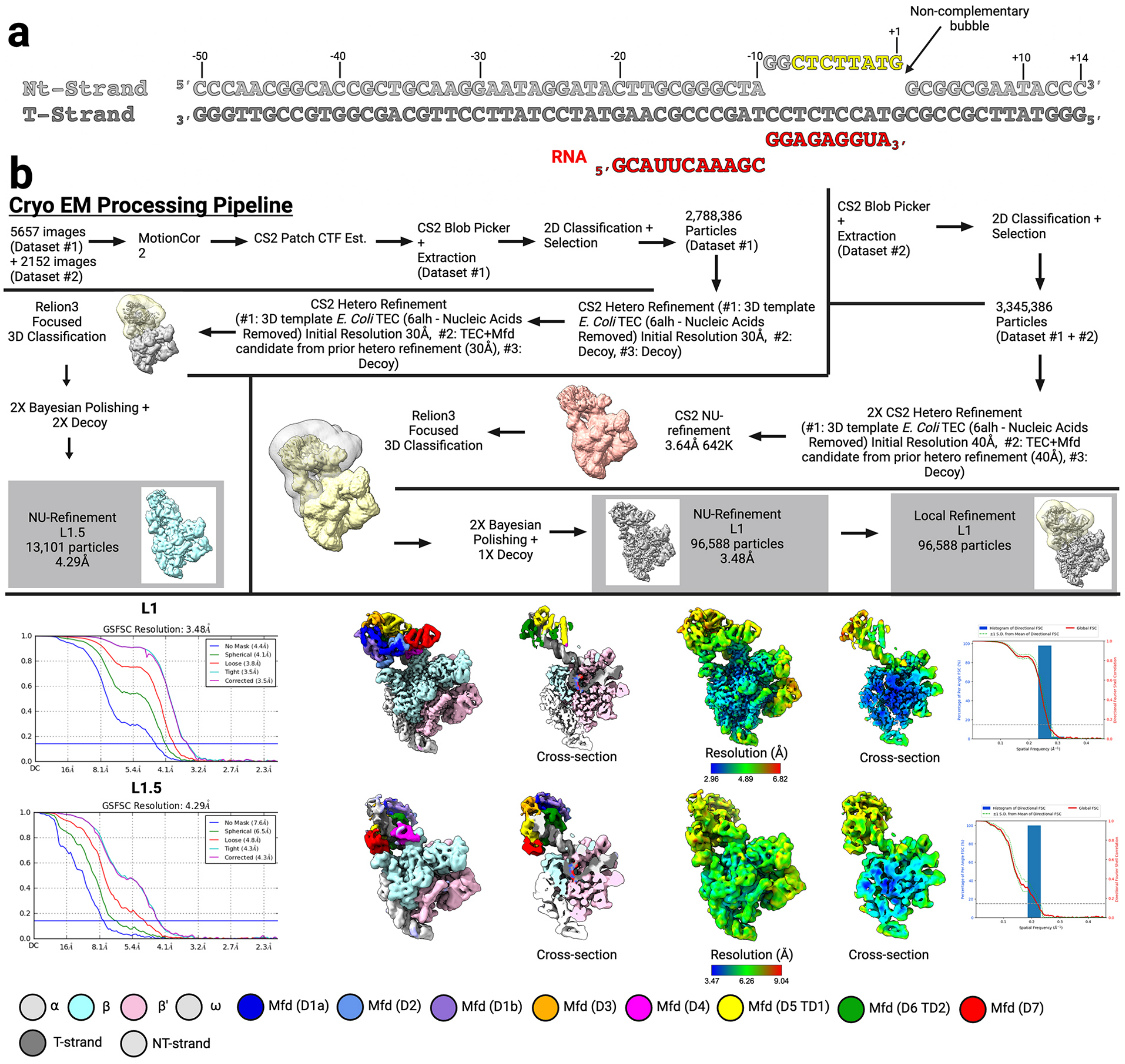

**Supplementary Figure 2.**

**(a)** Cryo-EM scaffold sequence highlighting partial complementarity between the RNA and t-strand DNA and between the t-strand and nt-strand.

**(b)** (Top half) Cryo-EM processing pipeline for L1 and L1.5 intermediates. (Bottom-half) Gold standard FSC calculations for cryo-EM density map [on left], cryo-EM density map and cross section colored according to key [middle-left], cryo-EM density map colored according to local resolution [middle-right] (Cardone et al. 2013), 3DFSC and sphericity of density map (Tan et al. 2017). L1 information shown in the upper row, L1.5 information shown in the lower row.

**Supplementary Table 1 | Cryo-EM data collection, refinement and validation statistics for Mfd-EC. Related to Figures 2&3.**

| **Dataset** | | |
|---|---|---|
| **Data collection and processing** | Dataset #1 | Dataset #2 |
| Microscope | FEI Titan Krios | FEI Titan Krios |
| Voltage (kV) | 300 | 300 |
| Detector | Gatan K3 | Gatan K3 |
| Electron exposure (e–/Å2) | 51.2 | 68.2 |
| Defocus range (μm) | 1 to 2.5 | 1 to 2.5 |
| Data collection mode | Counting Mode | Counting Mode |
| Pixel size (Å) | 1.0825 | 1.0825 (binned from 1.03) |
| Symmetry imposed | C1 | C1 |
| Initial particle images (no.) | 2,788,386 | 212,950 |
| **Refinement** | | |
| Structure | L1 | L1.5 |
| EMDB | EMD-48776 | EMD-48802 |
| PDB | 9N07 | 9N11 |
| Final particle images (no.) | 96,588 | 13,808 |
| Map resolution (Å) - FSC threshold 0.143 | 3.5 | 4.3 |
| Map resolution range (Å) | 3.0- 6.8 | 3.5- 9.0 |
| Initial model used (PDB code) | 6X26 | 6X50 |
| Map sharpening B factor (Å2) | 94.2 | 71.2 |
| Model composition | | |
| Non-hydrogen atoms | 71,263 | 70,701 |
| Protein residues | 4,316 | 4,261 |
| Nucleic acid residues (DNA/RNA) | 113 | 113 |
| Ligands | 2 $Zn^{2+}$, 1 $Mg^{2+}$, 1 ADP, 1 $BeF_3^-$ | 2 $Zn^{2+}$, 1 $Mg^{2+}$ |
| B factors (Å2) | | |
| Protein | 151.83 | 406.22 |
| Nucleic acid | 201.85 | 428.6 |
| Ligands | 283.02 | 279.08 |
| R.m.s. deviations | | |
| Bond lengths (Å) | 0.005 | 0.004 |
| Bond angles (°) | 0.606 | 0.635 |
| Validation | | |
| MolProbity score | 1.76 | 1.64 |
| Clashscore | 4.87 | 4.07 |
| Poor rotamers (%) | 0.17 | 1.0 |

| | | |
|---|---|---|
| Ramachandran plot | | |
| Favored (%) | 91.52 | 93.05 |
| Allowed (%) | 8.48 | 6.85 |
| Disallowed (%) | 0.0 | 0.09 |
